# Organization and Regulation of Chromatin by Liquid-Liquid Phase Separation

**DOI:** 10.1101/523662

**Authors:** B.A. Gibson, L.K. Doolittle, L.E. Jensen, N. Gamarra, S. Redding, M.K. Rosen

## Abstract

Genomic DNA is highly compacted in the nucleus of eukaryotic cells as a nucleoprotein assembly called chromatin^1^. The basic unit of chromatin is the nucleosome, where ∼146 base pair increments of the genome are wrapped and compacted around the core histone proteins^2,3^. Further genomic organization and compaction occur through higher order assembly of nucleosomes^4^. This organization regulates many nuclear processes, and is controlled in part by histone post-transtranslational modifications and chromatin-binding proteins. Mechanisms that regulate the assembly and compaction of the genome remain unclear^5,6^. Here we show that in the presence of physiologic concentrations of mono- and divalent salts, histone tail-driven interactions drive liquid-liquid phase separation (LLPS) of nucleosome arrays, resulting in substantial condensation. Phase separation of nucleosomal arrays is inhibited by histone acetylation, whereas histone H1 promotes phase separation, further compaction, and decreased dynamics within droplets, mirroring the relationship between these modulators and the accessibility of the genome in cells^7-10^. These results indicate that under physiologically relevant conditions, LLPS is an intrinsic behavior of the chromatin polymer, and suggest a model in which the condensed phase reflects a genomic “ground state” that can produce chromatin organization and compaction in vivo. The dynamic nature of this state could enable known modulators of chromatin structure, such as post-translational modifications and chromatin binding proteins, to act upon it and consequently control nuclear processes such as transcription and DNA repair. Our data suggest an important role for LLPS of chromatin in the organization of the eukaryotic genome.

---

It has been known for many years that various cations promote self-association of chromatin, resulting in its precipitation from solution^11,12^. Recent research has demonstrated that weak multi-valent interactions can cause LLPS of many biological molecules, producing highly dense liquid droplets^13,14^. Given polymer melt and fractal models of chromatin structure^15-17^, reported observations of cation-induced spherical aggregates of nucleosomal arrays^18^, and a view of chromatin as a highly-valent array of nucleosomes, we asked what the physical nature of cation-driven chromatin precipitates might be. We reconstituted nucleosome arrays composed of recombinant purified and fluorophore-labeled histone octamers and a defined DNA template containing 12 repeats of Widom’s 601 nucleosome positioning sequence (Figs. 1A and S1). Solutions of these purified arrays are homogeneous in the low salt conditions used at the end of the reconstitution protocol. However, confocal fluorescence microscopy revealed that addition of mono- or divalent cations to physiologically relevant concentrations resulted in formation of round, liquid droplets of phase-separated chromatin (Fig. 1B, D). These droplets were dependent on the assembly of DNA and histone octamers into nucleosomal arrays, and were not observed with either free DNA or histones, or with aggregates formed by addition of histones to DNA in physiologic salt (Fig. S2). Chromatin droplet formation did not require the addition of crowding agents and was independent of presence or type of fluorophore label, histone octamer species of origin, and treatment of the microscopy glass (Figs. 1C and S3). A characteristic trait of molecules that undergo phase separation is their sharp transition from a homogeneous solution to immiscible phases at defined threshold concentrations that depend on buffer conditions and are favored by higher molecular valency^19^. In this regard, we titrated monovalent (KOAc) and divalent (Mg[OAc]_2_) salts, and varied the number of nucleosomes in each array as well as array concentration. This revealed behaviour consistent with phase separation: chromatin droplets appeared sharply as salt or array concentrations were increased, and were favored by increasing nucleosome number in the arrays (Figs. 1D, E, and S4, 5). To ensure that phase separation of nucleosomal arrays was not a peculiarity of the assembly process, we designed an alternative approach wherein nucleosomal arrays were generated through DNA ligation of pre-assembled mononucleosomes (Figs. 1F, G). Ligation of mononucleosomes by T4 DNA ligase produced material with cation-dependent phase separation analogous to that of the dodecameric nucleosomal arrays (Figs. 1H, 1I).

**Figure 1.**
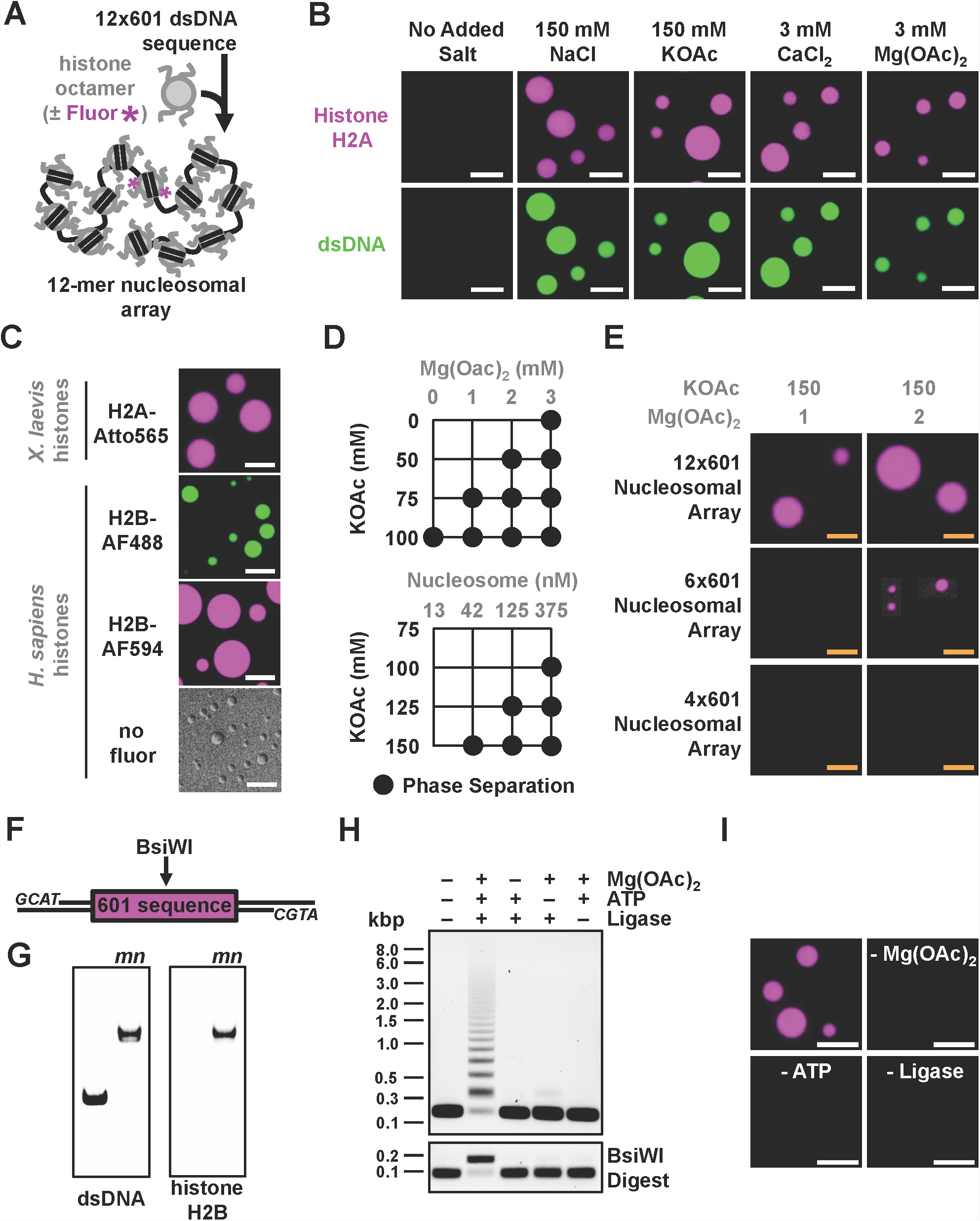
Phase separation of nucleosome arrays in physiologic salt. (A) Schematic depicting the assembly of a dodecameric nucleosome array from purified histone octamers, with or without a fluorophore (magenta) using a double stranded DNA (dsDNA) template encoded with repeats of Widom’s 601 nucleosome positioning sequence. (B) Confocal fluorescence microscopy images of Atto 565-labelled (via histone H2A) dodecameric nucleosome arrays and YOYO-1-labelled dsDNA, in magenta and green respectively, following addition of mono- or divalent cation salts. (C) Confocal fluorescence microscopy images of phase separated dodecameric nucleosome arrays assembled with *X. laevis* or *H. Sapiens* histone octamers labelled on histone H2A with Atto565, or histone H2B with Alexa Fluor 488 or Alexa Fluor 594 dyes (*top* to *middle bottom*). Differential interference contrast microscopy images of phase separated dodecameric nucleosome arrays assembled with *H. Sapiens* histone octamers without a conjugated dye (*bottom)*. (D) Phase diagrams determined by confocal microscopy in titrations of potassium acetate (KOAc) and either Mg(OAc)_2_ (top) or nucleosome array (bottom). (E) Confocal fluorescence microscopy images of AlexaFluor594-labelled arrays with different numbers of nucleosomes at identical total nucleosome concentration (100 nM). (F) Schematic depicting the molecular features of a DNA template for assembling mononucleosomes containing a 601 nucleosome positioning sequence, directionally ligate-able ends, and a BsiWI restriction site. (G) In-gel imaging of ethidium bromide-stained DNA separated by electrophoresis on a 6% native PAGE gel, with (mn lanes) and without assembly into mononucleosomes. Left gel shows ethidium bromide fluorescence, right gel shows Alexa Fluor 488-histone H2B fluorescence. (H) Following T4 DNA ligase-mediated joining of mononucleosomes, isolated ligation products with (bottom) and without (top) digestion by BsiWI were separated by agarose gel electrophoresis and visualized with ethidium bromide. (I) Confocal fluorescence imaging of AlexaFluor488-labeled chromatin droplets (upper left, false-colored in magenta). In the absence of magnesium (upper right), ATP (lower left) or T4 DNA ligase (lower right) ligation does not occur (panel H) and droplets do not form. Scale bars, in orange and white, are 4 and 10 µm, respectively.

Phase separated polymers can exhibit a variety of material properties, from rigid solids to dynamic liquid-like structures^20^. Chelation of free magnesium by super-stoichiometric addition of EDTA resulted in rapid dispersion of magnesium-dependent chromatin droplets (Fig. 2A), suggesting droplets can exchange cations readily with the surroundings. In contrast, however, photobleaching of the labeled histone in entire chromatin droplets resulted in very slow recovery of fluorescence (Figs. 2B, C), an observation more often found in solid-like phases^21,22^. We wondered if the slow fluorescence recovery was due, not to solid-like properties, but rather an absence of material in bulk solution from which to recover. Quantitation of nucleosome concentration in chromatin droplets (∼340 µM) and in bulk solution (∼30 nM) revealed a >10,000-fold concentration of nucleosomal arrays following LLPS, indicating that the inability to recover fluorescence of entire droplets following their photobleaching likely results from a dearth of free material in solution (Figs. 2D and S6). Short time-scale recovery of fluorescence following photobleaching of a portion of the chromatin droplets,confirmed the dynamic liquid-like properties of phase-separated chromatin (Fig. 2E). Internal droplet dynamics drive this fluorescence recovery, as fluorescence was lost outside of, and gained within, the irradiated volume following partial photobleaching of droplets at an equal and opposite initial rate (Fig. 2F). Given the very slow exchange of fluorescence with bulk solution, we could directly image droplet fusion and subsequent internal mixing of materials by co-incubating differentially labeled chromatin droplets. This revealed that fusion occurred rapidly, with droplets changing from an initial hourglass shape to spherical within ∼30 seconds. However, internal mixing was much slower, occurring on timescales of 10-20 minutes. Together, these behaviors indicate that chromatin droplets have high surface tension and high viscosity (Fig. 2G and Movie S1).

**Figure 2.**
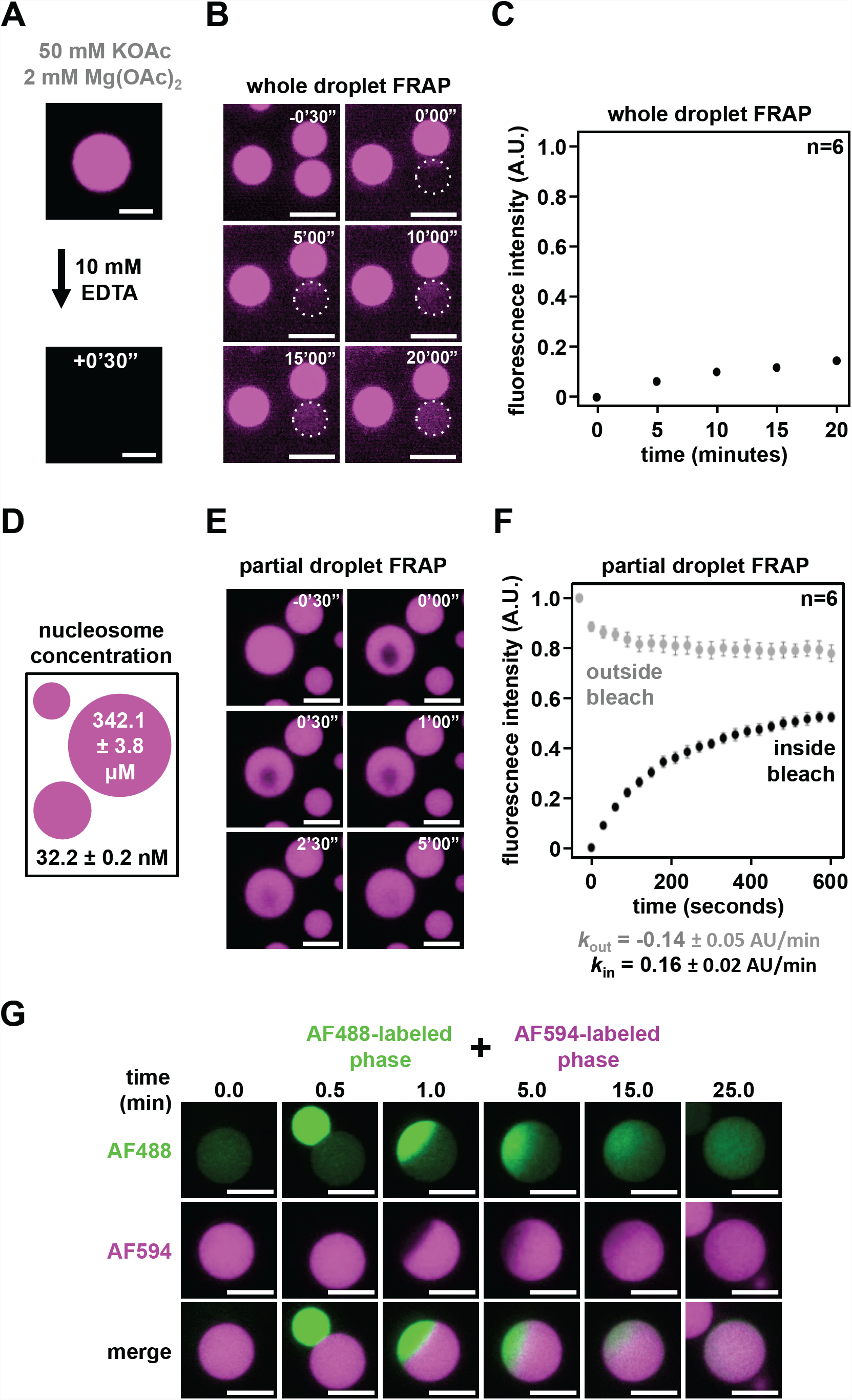
Chromatin droplets are highly concentrated and liquid-like. (A) Confocal fluorescence microscopy images of Alexa Fluor 594-labeled chromatin before and 30 seconds after addition of 10 mM EDTA. (B) Confocal fluorescence microscopy images of fluorescence recovery of whole Alexa Fluor 594-labeled chromatin droplets following photobleaching. (C) Quantification from 6 individual photobleaching experiments from panel B. (D) Nucleosome concentration within chromatin droplets and in the surrounding bulk solution. (E) Confocal fluorescence microscopy images of fluorescence recovery following partial photobleaching of Alexa Fluor 594-labeled chromatin droplets. (F) Quantification of fluorescence change within chromatin droplets both inside (black) and outside (grey) of the photobleached regions, averaged for 6 individual photobleaching experiments from panel E. Error bars and ± error are standard deviation. Initial rates of fluorescence change within chromatin droplets both inside (***k***_**in**_) and outside (***k***_**out**_) of the photbleached region are indicated in black and grey, respectively. (G) Confocal fluorescence microscopy images of fusion of Alexa Fluor 488- and Alexa Fluor 594- labeled chromatin droplets. The differently labeled droplets were formed separately and then mixed. Scale bars are10 µm.

The most abundant chromatin-binding protein in the majority of eukaryotes is the linker histone H1, which binds at the dyad axis of the nucleosome and plays roles in genomic access, gene regulation, and compaction in cells^23^. Given recent reports that the lysine-rich C-terminal tail of histone H1 can form coacervates with DNA^24^, we wondered how binding of histone H1 to nucleosomal arrays (Fig. 3A) might affect the phase separation of chromatin and material properties of the resulting droplets. Addition of purified calf thymus histone H1 to dodecameric nucleosomal arrays promoted phase separation of chromatin at half the concentration of monovalent salt compared to nucleosomal arrays alone (Fig. 3B). At equivalent concentrations, phase separated histone H1-bound chromatin droplets were generally smaller than those of chromatin alone, and had ∼1.5-fold higher fluorescence intensity independent of fluorophore labeling strategy (Figs. 3C, D, and S7). These data suggest a further compaction and concentration of phase-separated material as a result of histone H1 binding. Imaging of histone H1-bound chromatin droplets revealed the presence of unresolved fusion intermediates (Fig. 3B), and these droplets did not appreciably recover from partial photobleaching (Fig. 3E). These data indicate that histone H1 promotes phase separation of chromatin, increases compaction of nucleosomes within condensates, and decreases the dynamics of chromatin droplets. By implication, histone H1 may also exert a larger-scale decrease in chromatin dynamics in cells, perhaps explaining how H1 depletion in eukaryotic cells results in a loss of chromatin cohesion and increased nuclear volume^25^.

**Figure 3.**
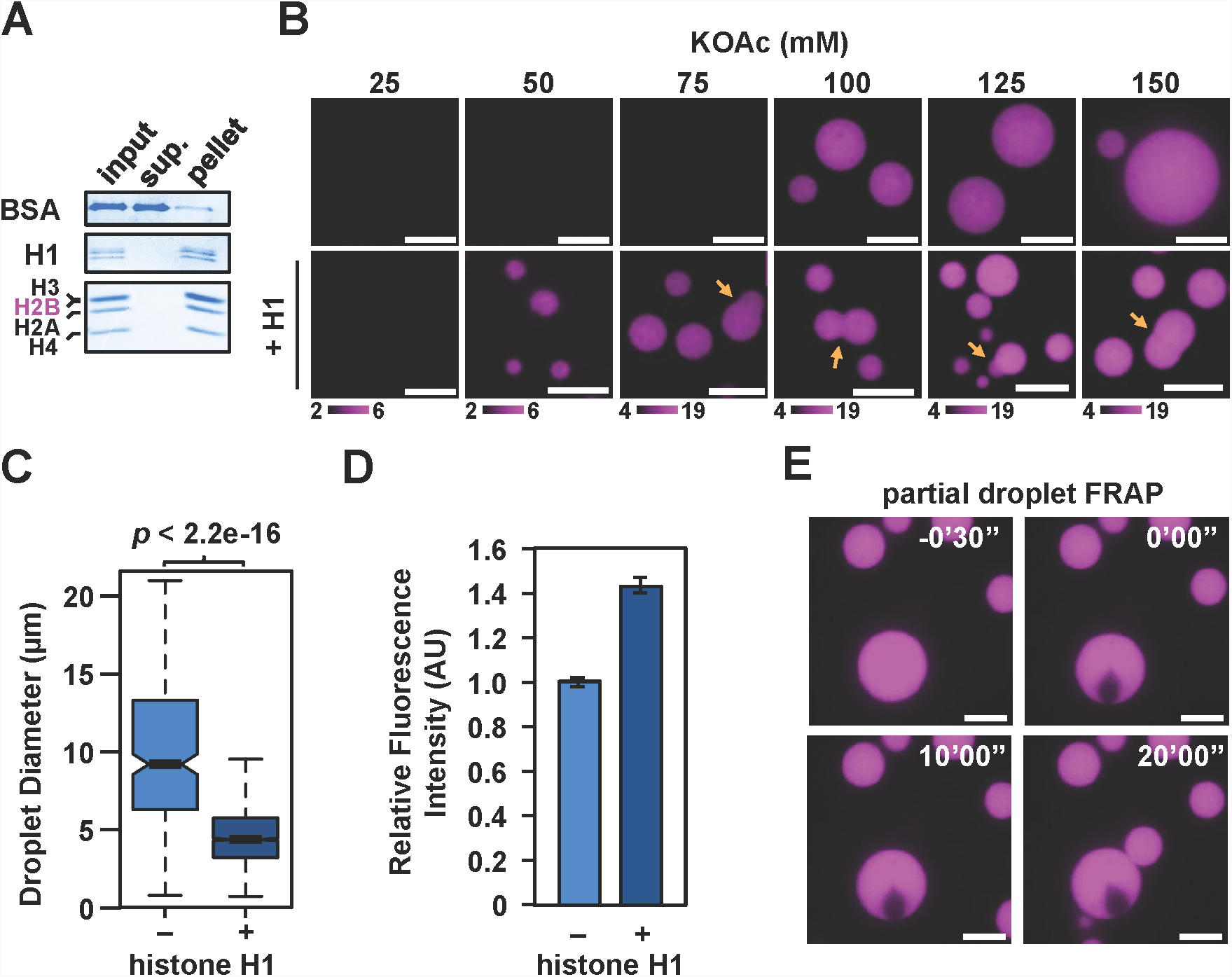
Histone H1 promotes phase separation of chromatin droplets with increased compaction and decreased dynamics. (A) Coomassie brilliant blue-stained SDS-PAGE gel of proteins in supernatant (sup.) or pellet following sedimentation of chromatin droplets containing histone H1. Histone H2B (magenta) is labeled with Alexa Fluor 594 and runs at the histone H3 position on a 15% PAGE-SDS gel. (B) Confocal fluorescence microscopy, with identical microscope settings and image processing in each buffering condition, of Alexa Fluor 594-labeled chromatin following titration of potassium acetate with (bottom) or without (top) histone H1. Fluorescence microscopy image pixel intensities are indicated below each buffering condition, with an enumerated color gradient from black to magenta. Orange arrows indicate stalled droplet fusion intermediates. (C) Quantification of droplet size from 3,336 droplets bound by histone H1 and 611 droplets not bound by histone H1. P-value was determined using the *t*-test. (D) Quantification of relative fluorescence intensity from 10 chromatin droplet centers either bound or not bound by histone H1. Error bars indicate standard deviation from the mean. (E) Confocal microscopy images of fluorescence recovery of Alexa Fluor 594-labeled chromatin in the presence of histone H1 following partial droplet photobleaching. Scale bars are 10 µm.

Nucleosomes associate with one another through a variety of mechanisms, including histone tail-DNA interactions and contacts between the “acidic” and “basic” patches of the nucleosome^26,27^. We asked whether these mechanisms also contribute to LLPS. Similar to previous observations of chromatin precipitation^28,29^, we found that nucleosomal arrays without histone tails, generated by partial proteolysis with trypsin (Fig. S8), do not undergo LLPS in the presence of physiologic salts (Fig. 4A). The histone H4 basic patch (H4K16, R17, R19, and K20) plays important roles in nucleosome array self-association through interactions with either DNA or the histone H2A acidic patch (H2A E61, E64, D90, and E92)^30^. We found that nucleosome arrays with charge-neutralizing mutations in the basic patch are defective in chromatin droplet formation, whereas acidic patch mutation does not appreciably inhibit phase separation (Fig. 4B), suggesting that interactions of the histone H4 basic patch with DNA are likely important determinants in chromatin LLPS.

**Figure 4.**
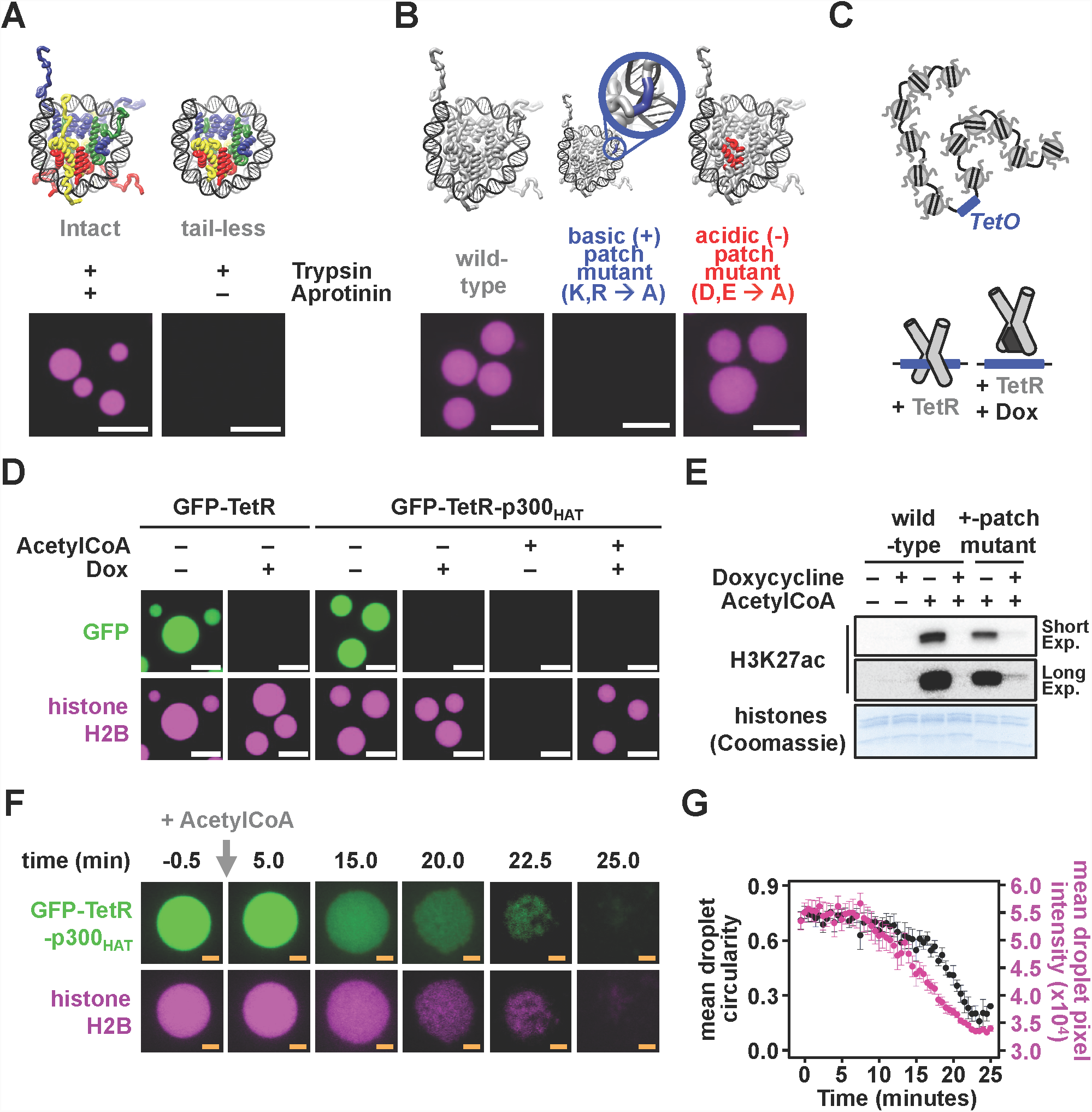
Histone acetylation dissolves chromatin droplets. (A) Confocal fluorescence microscopy images of Alexa Fluor 594-labelled chromatin following 30 minutes of trypsin-mediated digestion of histone tails in the presence or absence of trypsin inhibitor Aprotinin. (B) Confocal fluorescence microscopy images of fluorophore-labeled dodecameric nucleosome arrays at 375 nM nucleosome concentration in a buffered solution with 150 mM KOAc, assembled with wild-type, basic patch mutant, or acidic patch mutant *X. laevis* histone octamers. (C) A dodecameric nucleosome array containing a Tet Operator (*TetO*) can bind Tetracycline Repressor (TetR) in the absence of doxycycline (Dox). (D) Confocal fluorescence microscopy images of Alexa Fluor 594-labeled chromatin (magenta) and sfGFP fused to either the model transcription factor TetR (GFP-TetR) or TetR fused to the catalytic domain of p300 (GFP-TetR-p300_HAT_) (both green) including doxycycline (Dox) and/or AcetylCoA. (E) (top) Western blot of histone H3K27 acetylation following addition of doxycycline and/or AcetylCoA to droplets of *TetO*-containing dodecameric nucleosome arrays containing wild-type or basic (+)-patch mutant histones in the presence of GFP-TetR-p300_HAT_ and (bottom) Coomassie brilliant blue-stained SDS-PAGE gel of core histone proteins. (F) Confocal fluorescence microscopy of Alexa Fluor 594-labeled chromatin (magenta) and GFP-TetR-p300_HAT_ (green) following addition of AcetylCoA. (G) Mean droplet circularity and pixel intensity of Alexa Fluor 594-labeled chromatin droplets in the presence of GFP-TetR-p300_HAT_ following addition of AcetylCoA. Error bars indicate standard error. Scale bars, in orange and white, are 4 and 10 µm, respectively.

Nucleosome histone tails are acetylated *in vivo*, often by histone acetyltransferase enzymes recruited by transcription factors to specific loci, in order to regulate gene expression^7,31^. These modifications impair self-interaction and precipitation of nucleosomal arrays in vitro^32^, similar to basic patch mutations. To examine how acetylation might alter the formation and material properties of chromatin droplets, we devised a model system to mimic transcription factor-driven histone acetylation. We genetically linked the model *E. coli* transcription factor, Tet Repressor (TetR), to the catalytic domain of the histone acetyltransferase, p300 (p300_HAT_) and GFP, and combined this fusion protein (GFP-TetR-p300_HAT_) with a nucleosome array containing a central Tet Operator (*TetO*). In this system, the tetracycline analog, doxycycline (Dox), inhibits transcription factor binding to the chromatin, and Acetyl-CoA is necessary for histone acetylation by p300_HAT_ (Fig. 4C). GFP-TetR and GFP-TetR-p300_HAT_ were both strongly recruited to TetO-containing chromatin droplets, an effect that was blocked by Dox (Fig. 4D). Addition of Acetyl-CoA caused dissolution of chromatin droplets containing GFP-TetR-p300_HAT_, concomitant with acetylation of H3K27 and likely other sites as well (Fig. 4E). This effect required recruitment of GFP-TetR-p300_HAT_ into the droplets, as acetylation and droplet dissolution were blocked by Dox (Figs. 4D, E). By comparing GFP-TetR-p300_HAT_-mediated histone acetylation of phase separated wild type chromatin and non-phase separating basic patch mutant chromatin we found that wild type has both increased transcription factor-dependent acetylation (-Dox), and decreased transcription factor-independent acetylation (+Dox). Thus, compaction and LLPS enhance the fidelity of this signaling pathway (Fig. 4E). Time-resolved imaging of acetylation-mediated dissolution of chromatin droplets (Fig. 4F and Movie S2) shows that following an initial delay after the reaction is initiated, the density of droplets (assessed by fluorescence intensity) progressively decreases until the structures disappear. Droplets maintain their size and approximate shape through the early stages of this process until density (i.e.fluorescence intensity) decreases to roughly half its initial value, at which point they begin to crumple and lose circularity (Fig. 4G). These behaviors mimic transcription-factor-driven cellular signaling pathways, where localized histone acetylation stimulates decompaction of the genome in order to transactivate DNA-templated processes^31,33^. Moreover, they illustrate how acetylation can tune the density and material properties of chromatin droplets in vitro and likely in vivo as well.

Here we have demonstrated that in the presence of physiologic salt, the poly-nucleosome chromatin polymer has an intrinsic ability to form a highly compact, yet dynamic, liquid phase. Previous reports of self-associated chromatin aggregates^18,32,34^ can now be described as dynamic liquid-liquid phase separated droplets with unique viscoelastic material properties, that can be modulated by chromatin-binding proteins and signaling molecules in a manner consistent with their functions in cells. The density of this phase (∼340 µM nucleosome concentration) is similar to estimates of chromatin density in cells (∼80-520 µM)^35^, indicating that LLPS is sufficient produce the degree of compaction necessary to organize the genome in the nucleus. We have shown that charge neutralizing mutations of the histone H4 “basic patch” and hyperacetylation of histone lysines are incompatible with LLPS of chromatin. These data may explain the lethality of similar mutations in *S. cerevisiae*^*^36-38^*^ and *D. melanogaster*^*^39^*^ as well as the increased nuclear volume for both histone H4 lysine mutant survivors^37^ and cells with hyperacetylated chromatin following treatment with histone deacetylase inhibitors^40-42^. Although the droplets are dense, chromatin organized through LLPS can nevertheless be accessed and modulated by regulatory factors such as histone H1 and transcription factors (and very likely others), which would be necessary during nuclear processes such as transcription, replication and repair. However, as shown in our model transactivation assays, access can require recruitment of factors, suggesting that some molecules may be excluded from or inhibited by compaction within the dense phase. The droplets fuse rapidly, but the rate of content mixing is very slow, and should decrease further as the length of nucleosome arrays increases. Taken together with accessibility to regulatory factors, this behavior suggests that LLPS could allow regions of the genome to remain dynamic on shorter length scales while maintaining their spatial integrity on longer length scales within the nucleus. LLPS, as a mechanism to produce a compacted “ground state” of chromatin, could offer an avenue through which factors might produce a variety of “excited” structural states necessary for functional regulation of the genome.

## Acknowledgements

We thank Geeta Narlikar for advice and sharing unpublished results. Research was supported by the Howard Hughes Medical Institute and grants from the NIH (F32GM129925 to B.A.G.), the Welch Foundation (I-1544 to M.K.R.), and funding from the UCSF Program for Breakthrough Biomedical Research (PBBR) provided by the Sandler Foundation to S.R.

