## Supplementray Information for "Organization and Regulation of Chromatin by Liquid-Liquid Phase Separation"

**Gibson *et al.* (2018)**

This document contains the following supplementary information: Page

- Figure S3. Wetting of chromatin droplets with differential preparation of microscopy glass. . ........................................................................ 6
  - *Purification of H. sapiens Histones H3C111A, H4, and H2A Expressed in E. coli* .. 13
    - Expression ........................................................................ 13
    - Purification ........................................................................ 13
  - *Purification of Fluorophore-labeled H. sapiens Histone H2B Expressed in E. coli*. 14

### 1) Supplementary Movies and Figures

**Movie S1. Imaging of two-color Droplet-droplet Fusion.**
Following separate phase separation, 1% AF488-labeled dodecameric nucleosomal arrays (in green) and 1% AF594-labeled dodecameric nucleosomal arrays (in magenta) were mixed in a microscopy well and confocal fluorescence microscopy images were captured every 15 seconds at a using a spinning disk confocal microscope.

*[See MovieS1.mp4 file included with this paper]*

**Movie S2. Histone acetylation-dependent dissolution of chromatin droplets.**
Following in-phase binding of GFP-TetR-p300$_{HAT}$ (in green) to 1% AF594-labeled chromatin droplets (in magenta) composed of 12x601 *TetO*-containing nucleosomal arrays, AcetylCoA was added to the microscopy well, between frames 1 and 2, and mixing at 400 μM final concentration achieved by diffusion. Confocal fluorescence microscopy images were captured every 15 seconds at a using a spinning disk confocal microscope.

*[See MovieS2.mp4 file included with this paper]*

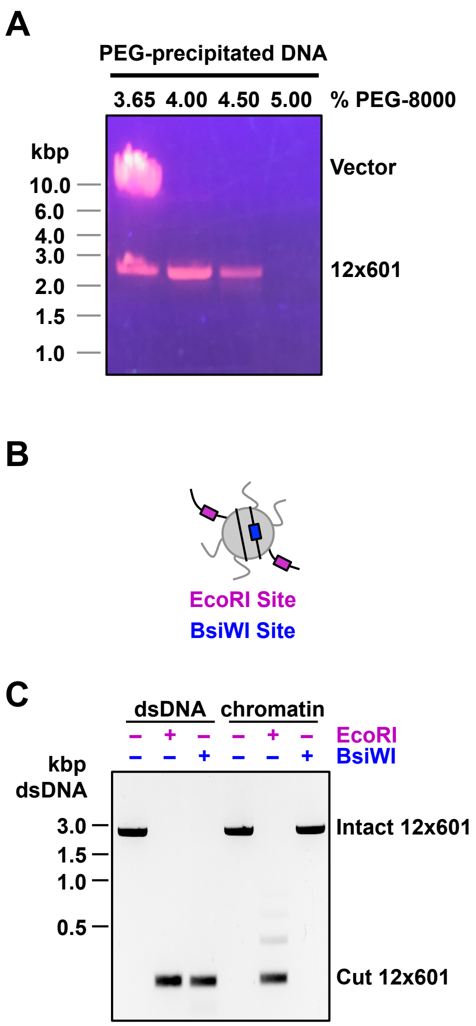

**Fig. S1. 601 DNA array preparation and differential digest of nucleosomal arrays.**
(A) Large scale PEG-fractionated DNA isolated from Phenol-chloroform extracted EcoRV-HF digestion products of the p21x601 plasmid run separated on a 1% agarose gel in 1xTAE stained with ethidium bromide. (B) Diagram depticitng EcoRI and BsiWI restriction endonuclease recognition sites relative to a well-positioned nucleosome within the 12x601 nucleosomal array. (C) Differential digestion of naked 12x601 dsDNA and 12x601 DNA assembled into a nucleosomal array using EcoRI-HF and BsiWI-HF restriction endonucleases. Cut DNA was extracted using a Qiagen PCR Purification Kit and run on a 1% Agarose Gel in 1xTAE.

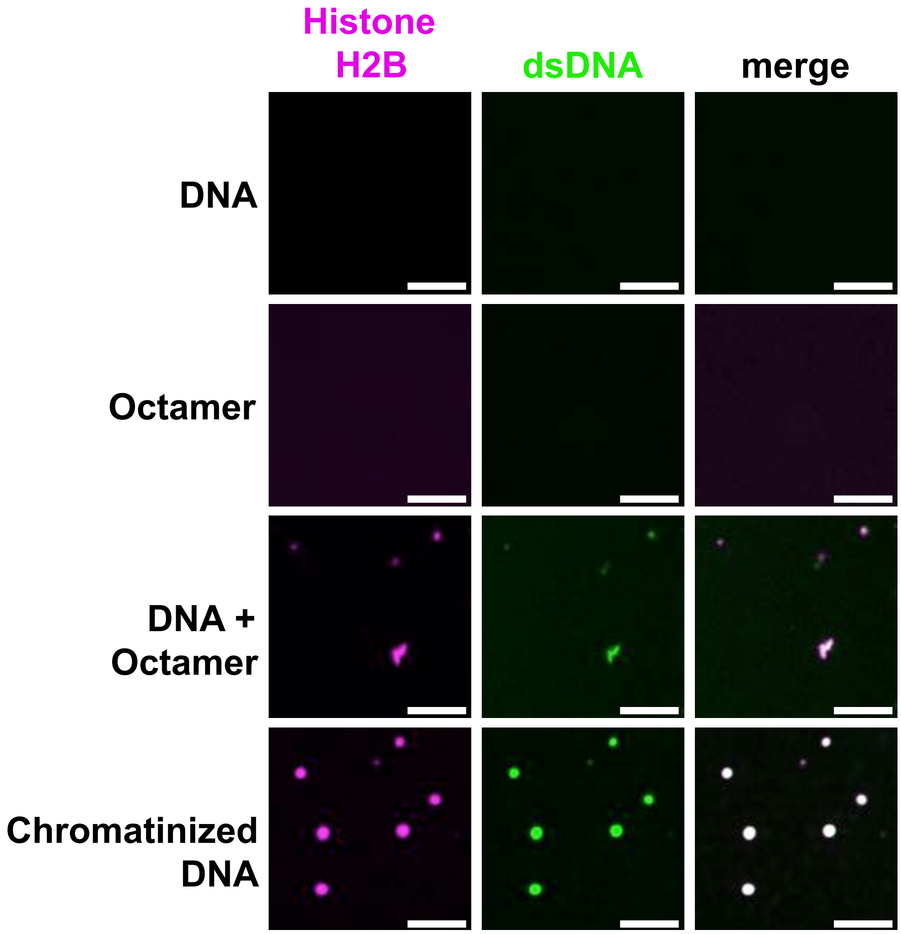

**Fig. S2. Assembled nucleosome-dependent formation of chromatin droplets.**
Confocal fluorescence microscopy images of 12x601 DNA alone, histone octamer alone, free histones mixed with 12x601 DNA, and nucleosomal arrays prepared by salt dialysis-mediate assembly. DNA, in green, was labeled with YOYO-1 and histones, in magenta, are labeled on histone H2B with AF594. Scale bars are 10 µm.

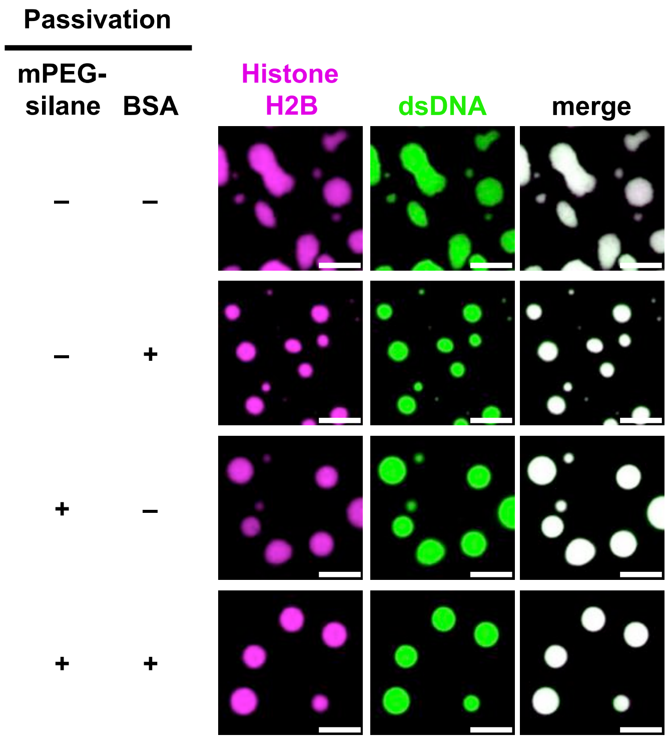

**Fig. S3. Wetting of chromatin droplets with differential preparation of microscopy glass.** Confocal fluorescence microscopy images of chromatin droplets labeled on histone H2B with AF594 and with dsDNA labeled using YOYO-1. Droplets were added to the well of a microscopy plate treated with and without mPEGylation and with and without passivation with BSA. Scale bars are 10 μm.

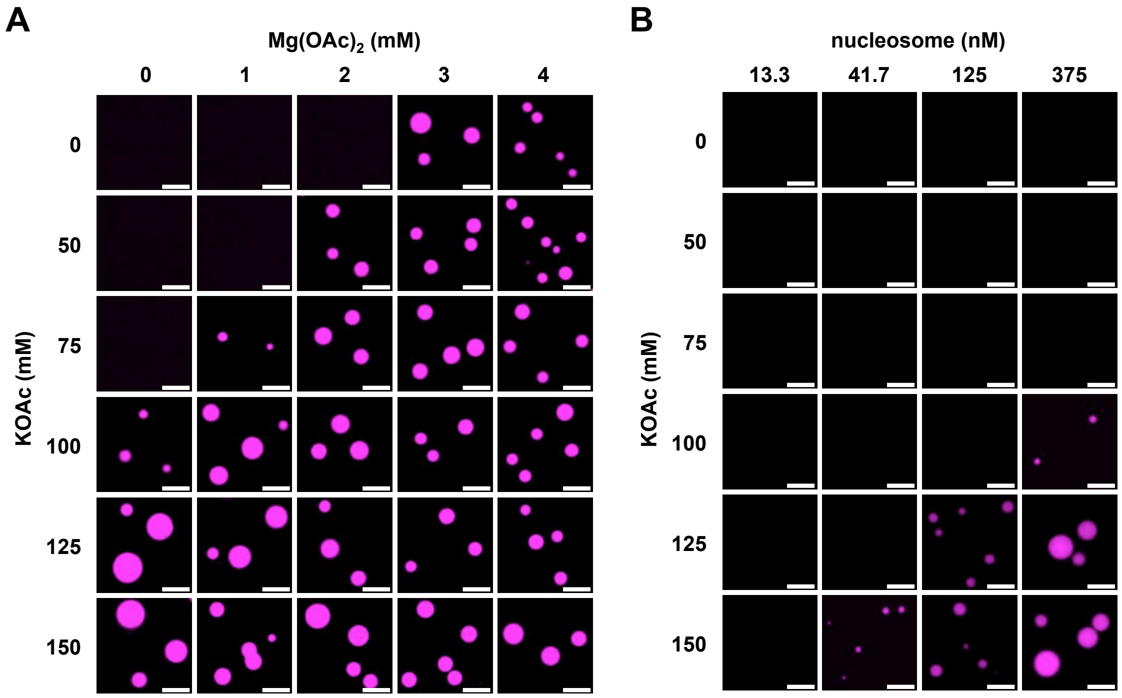

**Fig. S4. Modulation of chromatin droplet formation by titration of monovalent salt, divalent salt, and nucleosome concentrations.**
Confocal fluorescence microscopy images of dodecameric nucleosomal arrays assembled with *X. laevis* histone octamers labelled on histone H2A with Atto565 following titration of (A) KOAc and $Mg(OAc)_2$ at or (B) KOAc and nucleosomal arrays. Scale bars are 10 µm.

.

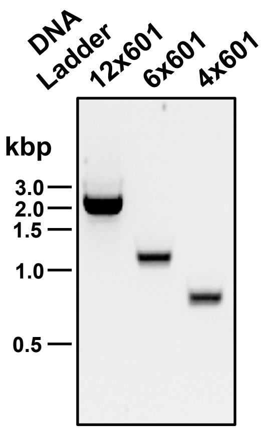

**Fig. S5. 12x601, 6x601, and 4x601 Nucleosomal Array Templates .**
1% agarose gel electrophoresis of 12x601, SacI-digested 6x601, and KpnI- and NcoI-digested 4x601 dsDNA with staining by ethidium bromide.

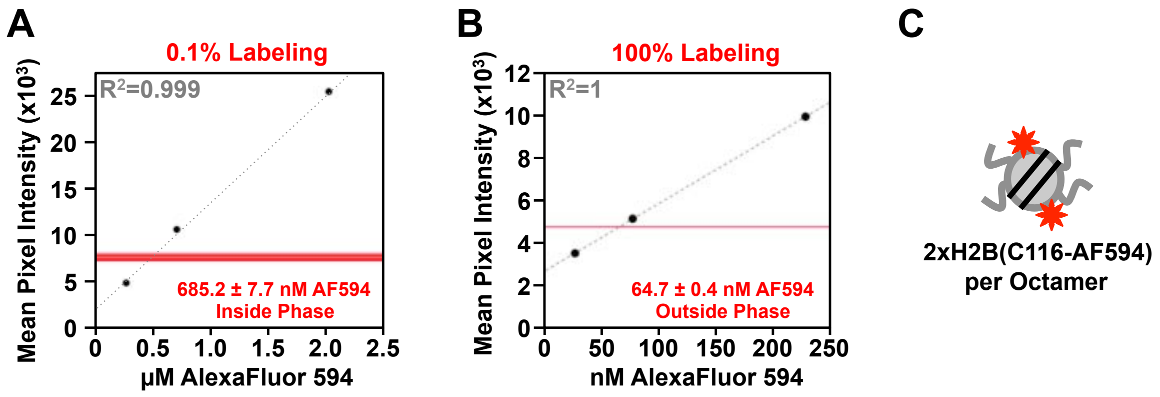

**Fig. S6. Quantitation of nucleosome concentrations both within and without chromatin droplets.**

Quantitation of chromatin droplets from confocal fluorescence microscope images. (A) Mean droplet intensity of chromatin droplets composed of dodecameric nucleosomal arrays with 0.1% AF594 doubly-labeled histone octamers relative to a standard curve of free AF594 dye. Mean droplet intensity of 10 individual chromatin droplets are depicted with horizontal red lines. (B) Three independent experiments quantitating fluorophore content in supernatant above pelleted chromatin condensates composed of dodecamieric nucleosomal arrays prepared with nucleosomes 100% doubly-labeled with Alexa Fluor 594 relative to a standard curve of free Alexa Fluor 594 fluorescent dye (filled black circle). (C) Diagram depicting labelling scheme of histone octamers highlighting 2 dyes per octamer.

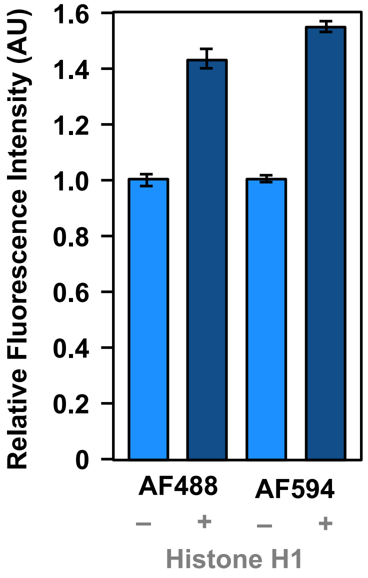

**Fig. S7. Histone H1 increases droplet fluorescence independent of fluorophore.**
Bar graph representation of relative mean fluorophore intensity of histone H1-bound and unbound chromatin droplets labeled with of 0.1% Alexa Fluor 488 (AF488) or 0.1% Alexa Fluor 594 (AF594).

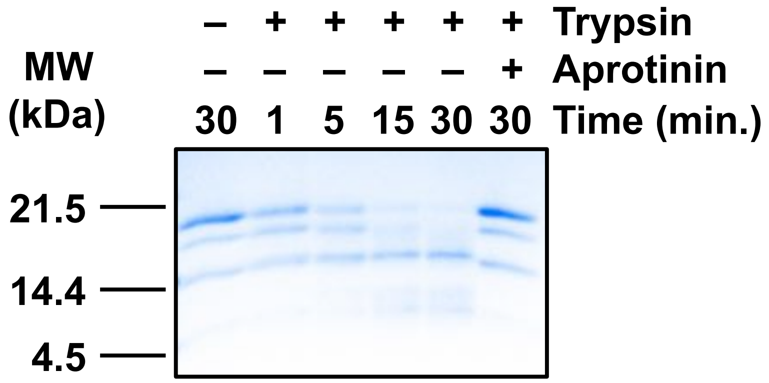

**Fig. S8. Trypsin-dependent removal of histone tails from nucleosomal arrays.**
Coomassie Brilliant Blue-stained 15% PAGE-SDS gel following proteolysis of dodecameric nucleosomal arrays with Trypsin.

### 2) Materials and Methods

#### Chemicals and Reagents

All chemicals and reagents were ordered at the highest possible purity available.

#### Molecular Biology and Cloning

***Construction of Bacterial Protein Expression Vectors.*** The following expression vectors were constructed for expression of the core histone proteins, the p300 catalytic domain, and fusion proteins GFP-TetR, and GFP-TetR-p300$_{HAT}$.

*H. sapiens Core Histones:* Synthetic open reading frames (ORFs) encoding *H. sapiens* histone H3C111A and H2BT116C were amplified from a dsDNA synthesized by Integrated DNA Technologies (IDT) using polymerase chain reaction (PCR) and primers adding a 5'-proximal NcoI restriction endonuclease recognition site and a 3'-proximal stop codon and BamHI restriction endonuclease recognition site. NcoI and BamHI restriction endonucleases from New England Biolabs (NEB) were used to directionally clone the H3C111A and H2BT116C ORFs into pET19b (Novagen). The sequence content of the protein expression vectors pET19b_H3C111A and pET19b_H2BT116C were confirmed by Sanger sequencing.

*X. laveis Core Histones:* pET-based protein expression constructs for the expression of wild-type and H3T33C and H2AK120C histone proteins from *X. laevis* were a generous gift from Dr. Geeta Narlikar. pET-based protein expression constructs for the expression of acidic patch mutant *X. laevis* histone H2A (H2A E61A, E64A, D90A, and E92A) and basic patch mutant X. laevis histone H4 (H4 K16A, R17A, R19A, K20A) were a generous gift from Dr. Song Tan.

*ySIR2:* A synthetic ORF encoding the histone deacetylase domain of *S. cerevisiae* protein SIR2 was amplified from a dsDNA synthesized by IDT using PCR and primers adding a 5'-proximal translation start codon and a 3'-proximal translation stop codon and XhoI restriction endonuclease recognition site. XhoI restriction endonuclease (NEB) digested PCR product encoding amino acids 87-562 of wild-type SIR2 was cloned into the pETduet-1 expression vector (Novagen) using a XhoI and a blunted NdeI restriction endonuclease digestion site (NEB). The sequence content of the protein expression vector pETduet_ySIR2 was confirmed by sanger sequencing.

*GFP-TetR and GFP-TetR-p300$_{HAT}$:* GFP-TetR and GFP-TetR-p300$_{HAT}$ were amplified using a two-step PCR method from (1) pEB1-sfGFP (Addgene #103983), (2) a synthetic sequence encoding *E. coli* TetR synthesized by IDT, and (3) a natural sequence encoding for amino acids 649-1026 of *H. sapiens* p300 (p300$_{HAT}$) synthesized by IDT (*1*). Primers used for PCR amplification of ORFs encoding the GFP-TetR and GFP-TetR-p300$_{HAT}$ fusion proteins added (1) a 5'-proximal BamHI restriction endonuclease recognition site, (2) a 3'-proximal translation stop codon followed by a NotI restriction endonuclease recognition site, and (3) sequence encoding for flexible $(GGS)_3$ linkers in between the domains of each fusion protein. BamHI-HF and NotI-HF restriction endonucleases (NEB) were used to directionally clone restriction endonuclease digested PCR products into the pETduet_ySIR2 expression vector (New England Biolabs) to encode for GFP-TetR and GFP-TetR-p300$_{HAT}$ with an N-terminal HHHHHHHHHHENLYFQGS sequence. The sequence content of the protein expression vectors pETduet_GFP-TetR and pETduet_GFP-TetR-p300$_{HAT}$ were confirmed by Sanger sequencing.

***Construction of 12x601 dsDNA Array-Producing Bacterial Vector.*** The p601 plasmid(*2*), containing a 12x601 array with Tet Operator (TetO) inserted between 601 sequences 6 and 7 cloned in a pCR-Blunt II-TOPO vector background was modified by (1) adding an EcoRV site 3'-proximal to the 12x601 sequence and (2) adding a 3,697 bp SpeI-released fragment from pMD2.G 5'-proximal to the 12x601 sequence using an SpeI restriction endonuclease recognition site.

**Expression and Purification of Recombinant Proteins**

***Purification of X. laeviss Core Histone Proteins Expressed in E. coli.***

Expression: Recombinant histones from *X. Laevis* were expressed in E. coli as previously described (*3*).

Purification: Recombinant histones from *X. Laevis* were purified in E. coli as previously described (*3*), with some modification. Briefly, washed inclusion bodies containing *E. coli*-expressed histone proteins were solubilized with XL Unfolding Buffer (20 mM Tris•HCl, pH 7.5, 7 M Guanidinium•HCl, 5 mM ß-Mercaptoethanol) and dialyzed into XL Dialysis Buffer (20 mM Tris•HCl, pH 7.5, 6 M Urea, 5 mM ß-Mercaptoethanol). Histone proteins were purified from soluble dialysate in XL Dialysis buffer by denaturing cation exchange chromatography using a TSKgelSP-5PW (TOROH) column, eluting histone proteins with a linear gradient of 0-1 M NaCl. Fractions containing histones were dialyzed into >18 MΩ $H_2O$ and lyophilized prior to histone octamer reconstitution.

***Purification of H. sapiens Histones H3C111A, H4, and H2A Expressed in E. coli.***

Expression: An overnight culture of Rosetta 2 (pLysS) *E. coli* (Novagen) transformed with pET19b_H3C111A, pET28a_H4 (Addgene #42633), or pET28A_H2A.1 (Addgene #42634) plasmids encoding wild-type *H. sapiens* histones H4 or H2A or mutant *H. sapiens* histones H3C111A, were grown on an agar plate by re-plating a single transformant on LB supplemented with 100 ng/µL of ampicillin (pET19b_H3C111A) or 35 µg/mL Kanamycin (pET28a-based expression) and 25 ng/µL of chloramphenicol at 37°C (*4*). The bacterial lawn was suspended in LB supplemented with the appropriate antibiotics, as above, and grown to a density ($OD_{600\ nm}$) of 0.4. Recombinant protein expression was then induced by addition of IPTG to 1 mM for 3 hours at 37°C. The cells were collected by centrifugation, and the bacterial cell pellets were resuspended in Histone Lysis Buffer (50 mM Tris•HCl, pH 8, 150 mM NaCl, 5 mM ß-mercaptoethanol, 1 mM Benzamidine, 100 µM Leupeptin, 100 µM Antipain, 1 µM Pepstatin), flash frozen in liquid $N_2$, and stored at -80°C.

Purification: Histones were purified essentially as previously described ((*3*)), with some modifications. *E. coli* expressing histone H4, H2A, or H3C111A resuspended in Histone Lysis Buffer were thawed on wet ice and lysed by multiple passages through an Avestin Emulsiflex-C5 high pressure homogenizer at ~10,000 PSI. Histone-containing inclusion bodies were separated from soluble bacterial lysate by centrifugation in a Beckman Avanti J-26 XPI centrifuge in a JA25.5 rotor at 19,500 RPM. Soluble bacterial lysate was discarded, and inclusion bodies were washed by resuspension and in 25 mL of Inclusion Body Wash Buffer (50 mM Tris•HCl, pH 7.5, 100 mM NaCl, 1% Triton X-100, 1 mM EDTA, 1 mM Benzamidine, 5 mM ß-mercaptoethanol) per liter of bacterial expression followed by pelleting by centrifugation in a Beckman Avanti J-26 XPI centrifuge in a JA25.5 rotor at 19,500 RPM. Inclusion bodies were washed once more with Inclusion Body Wash Buffer and twice more with Inclusion Body Wash Buffer omitting Triton X-100. Inclusion Bodies were soaked for 30 minutes with 167µL DMSO per liter bacterial expression, minced with a spatula, and proteins were extracted for 1 hour by stirring in 5 mL of Histone Unfolding Buffer (20 mM Tris•HCl, pH 7.5, 7M Guanidinium-HCl, 10 mM DTT) per liter of bacterial expression. Extracted unfolded proteins were separated from precipitate by centrifugation in a Beckman Avanti 6 XPI centrifuge in a JA25.5 rotor at 19,500 RPM. Inclusion body proteins were extracted once more for 40 minutes with stirring in 1.7 mL of Histone Unfolding Buffer per liter bacterial expression. Extracted unfolded proteins were separated once more from precipitate by centrifugation in a Beckman Avanti J-26 XPI centrifuge in a JA25.5 rotor at 19,500 RPM.

Pooled unfolded soluble inclusion body proteins were filtered through a 0.45 µm membrane (GE Healthcare) and run in Histone Unfolding Buffer over a HiLoad 26/60 Superdex 200 pg size exclusion column. Fractions containing histone proteins were dialyzed twice against 5 mM ß-mercaptoethanol in >18 MΩ $H_2O$ and once in 50% Glycerol. Precipitated proteins were pelleted by centrifugation in a Beckman Avanti J-26 centrifuge in a JA25.5 rotor at 19,500 RPM. Histone proteins in Histone Storage Buffer (50% Glycerol, 5 mM ß-mercaptoethanol) were concentrated in centrifugal concentrators (Amicon) with a 3,000 dalton molecular weight cutoff (MWCO) and stored at -20˚C.

8 M Urea was deionized by 3 passages through Amberlite MB-20 resin (SIGMA) and used to make SAU200 (20 mM NaOAc, pH 5.2, 7 M Urea, 200 mM NaCl, 1 mM EDTA, 5 mM ß-mercaptoethanol) and SAU600 (20 mM NaOAc, pH 5.2, 7 M Urea, 600 mM NaCl, 1 mM EDTA, 5 mM ß-mercaptoethanol) Buffer. Histone proteins in Histone Storage Buffer stored at -20˚C were diluted in >20 volumes of SAU200, filtered through a 0.45 µm membrane (GE Healthcare), and applied to Source 15S Chromatography Resin (GE Healthcare) equilibrated in SAU200. After washing with SAU200 and a 5 column volume gradient of 0-30% SAU600, histone proteins were eluted over a 35 column volume gradient of 30-100% SAU600. Fractions containing histone proteins were pooled and dialyzed three times against 5 mM ß-mercaptoethanol in >18 MΩ $H_2O$. Histone proteins were concentrated in centrifugal concentrators with a 3,000 dalton MWCO to less than 500 µL per liter bacterial expression and concentration quantified by measuring histone protein absorbance at 280 nm and the calculated molar extinction coefficients (https://web.expasy.org/protparam/) for histones H3C111A, H4, and H2A of 4470/M•cm, 5960/M•cm, and 4470/M•cm, respectively. Purified histone proteins were aliquoted in 200 (histones H3C111A or H4) or 240 nmol (histone H2A) quantities, flash frozen with liquid $N_2$, and stored at -80 ˚C.

***Purification of Fluorophore-labeled H. sapiens Histone H2B Expressed in E. coli.***

Expression: An overnight culture of Rosetta 2 (pLysS) *E. coli* (Novagen) transformed with pET19b_H2BT116C plasmids encoding *H. sapiens* histone H2B with an exogeneously introduced cysteine at threonine position 116, were grown on an agar plate by re-plating a single transformant on LB supplemented with 100 ng/µL of ampicillin and 25 ng/µL of chloramphenicol at 37°C. The bacterial lawn was suspended in LB supplemented antibiotics, as above, and grown to a density ($OD_{600\ nm}$) of 0.4. Recombinant protein expression was then induced by addition of IPTG to 1 mM for 3 hours at 37°C. The cells were collected by centrifugation, and the bacterial cell pellets were resuspended in Histone Lysis Buffer (50 mM Tris•HCl, pH 8, 150 mM NaCl, 5 mM ß-mercaptoethanol, 1 mM Benzamidine, 100 µM Leupeptin, 100 µM Antipain, 1 µM Pepstatin), flash frozen in liquid $N_2$, and stored at -80°C.

Purification: Histone H2B was purified similar to previously described (*5*), with modification. *E. coli* expressing histone H2BT116C resuspended in Histone Lysis Buffer were thawed on wet ice and lysed by multiple passages through an Avestin Emulsiflex-C5 high pressure homogenizer at ~10,000 PSI. Soluble bacterial lysate was isolated by centrifugation of cellular debris in a Beckman Avanti J-26 XPI centrifuge in a JA25.5 rotor at 19,500 RPM. Urea was added to soluble bacterial lysate until achieving a concentration of 7 M, and 3 M NaOAc at pH 5.2 was added to lysate until the lysate was acidified to ~ pH 5.2. Lysate was filtered by passage through a 0.45 μm membrane (GE Healthcare), and applied to Source 15S Chromatography Resin (GE Healthcare) equilibrated in SAU200. After washing with SAU200 and a 5 column volume gradient of 0-30% SAU600, histone H2BT116C was eluted over a 35 column volume gradient of 30-100% SAU600. Fractions containing histone proteins were pooled and dialyzed twice against 5 mM ß-mercaptoethanol in >18 MΩ $H_2O$ and once against Histone Storage Buffer. Histone H2BT116C was concentrated in a centrifugal concentrator with a 3,000 dalton MWCO quantified by measuring histone protein absorbance at 280 nm and the calculated molar extinction coefficient (https://web.expasy.org/protparam/) for histones H2BT116C of 7450/M•cm. Purified histone H2BT116C was stored prior to labeling at -20˚C in Histone Storage Buffer.

*Labeling:* Tris-neutralized TCEP (500 mM Tris•HCl, pH 8, 100 mM TCEP) was added to 1 mM final concentration to histone H2BT116C in Histone Storage Buffer and incubated at room temperature for 1 hour. Histone H2BT116C with fully reduced cysteines was moved into Phosphate Buffered Saline (8 mM $Na_2HPO_4$, 2 mM $KH_2PO_4$, pH 7.4, 137 mM NaCl, 2.7 mM KCl) using 2x5 mL HiTrap Desalting Columns (GE Healthcare) and labeled by addition of 1.5 molar excess Alexa Fluor 488 (AF488)-$C_5$-maleimide or Alexa Fluor 594 (AF594)-$C_5$-maleimide followed by incubation in the dark for 4 hours at room temperature. Addition of DMSO alone was used to generate unlabeled H2BT116C protein. Conjugation reactions were quenched by addition of 10 mM DTT. Removal of free fluorophore and any free dsDNA was achieved by flowing unlabeled, AF488-labeled, or AF594-labeled histone H2B through 2x5mL desalting columns and Source 15Q resin (GE Healthcare) equilibrated in Histone CleanUp Buffer (20 mM Tris•HCl, pH 7.5, 150 mM NaCl, 1 mM DTT). Fractions containing unlabeled, AF488-labeled, or AF594-labeled histone H2B proteins were pooled and dialyzed three times against 5 mM ß-mercaptoethanol in >18 MΩ $H_2O$. Differential labeled proteins were concentrated in centrifugal concentrators with a 3,000 dalton MWCO to less than 500 μL per liter bacterial expression and both concentration and percent labeling were quantified by measuring absorbance at 280, 495, and 590 as well as absorbance at and the calculated molar extinction coefficients (https://web.expasy.org/protparam/ or Thermo Scientific) for histone H2BT116C, AF488, and AF594 of 4470/M•cm, 73000/M•cm, and 92000/M•cm, respectively. 100% labeling was confirmed, and the purified and differentially labeled histone H2B proteins were aliquoted in 240 nmol quantities, flash frozen with liquid $N_2$, and stored at -80 ˚C.

***Fluorophore Labeling of X. laevis Histones H2A and H3 Expressed in E. coli.***

Lyophilized purified X. laevis histone proteins H2A and H3 with engineered cysteines at positions 120 and 33, respectively, were resuspended in XL Labeling Buffer (20 mM Tris•HCl, pH 7.5, 6 M Guanidinium•HCl, 5 mM EDTA, 0.7 mM TCEP) and Atto565-$C_5$-maleimide (SIGMA) or Cy5-$C_5$-maleimide (GE) were added at a 1:5 molar ratio of histone protein: maleimide dye. Conjugation reactions were carried out for 12 hours at room temperature in the dark. Free unconjugated dye was removed from labeled histone proteins with a 5 mL HiTrap Desalting Column (GE) followed by dialysis into water. After percent conjugation and protein concentration of dialyzed histone proteins were quantified, histones were aliquoted and lyophilized for histone octamer reconstitution.

***Purification of Calf Thymus Histone H1.***

Calf thymus histone H1 (14-155; EMD Millipore) was diluted in 20 volumes of Histone H1 Buffer A (25 mM $K^{+}PO_4^{-}$, pH 6.8, 100 mM NaCl, 1 mM DTT) and flowed through a Source 15Q column to remove any free DNA and applied to a Source 15S column equilibrated in Histone H1 Buffer A. The Source 15S column was washed with 2 column volumes of Histone H1 Buffer A, and 0-20% Histone H1 Buffer B over 5 column volumes. Fractions containing histone H1 from a linear elution gradient of 20-100% Histone H1 Buffer B were concentrated with a 3,000 Dalton MWCO centrifugal concentrator and applied to a Superdex 75 Increase 10/300 GL column (GE Healthcare) equilibrated in Histone H1 Buffer A. Fractions containing purified histone H1 were concentrated with a 3,000 Dalton MWCO centrifugal concentrator and quantified by measuring histone H1 protein absorbance at 280 nm and the calculated molar extinction coefficient (https://web.expasy.org/protparam/) for histone H1.5 of 1490/M•cm. Purified histone H1 was aliquoted in single use quantities, flash frozen with liquid $N_2$, and stored at -80 ˚C.

***Purification of GFP-TetR and GFP-TetR-p300$_{HAT}$ Expressed in E. coli.***

*Expression:* Rosetta 2 (pLysS) *E. coli* (Novagen) were transformed with either pETduet_GFP-TetR or pETduet_GFP-TetR-p300$_{HAT}$ plasmids encoding a fusion protein composed of superfolder GFP, the *E. coli* Tetracycline Receptor (TetR), and for GFP-TetR-p300 the histone acetyltransferase domain of *H. sapiens* p300 with an N-terminal Tobacco Etch Virus protease cleavable 10xHIS tag and grown overnight on an agar plate with LB medium supplemented with 100 ng/µL Ampicillin and 25 ng/µL Chloramphenicol. A single colony was used to inoculate a liquid preculture of LB with Ampicillin and Chloramphenicol, as above, with growth for ~8 hours at 37˚C. The liquid preculture was centrifuged at 4000 RPM for 10 minutes in a benchtop centrifuge and resuspended in a small volume of LB medium to inoculate 3 L of Terrific Broth (Difco) containing 100 ng/µL Ampicillin and 25 ng/µL of Chloramphenicol with growth position 116, were grown on an agar plate by re-plating a single transformant on LB supplemented with Antifoam 204, 100 ng/µL of ampicillin, and 25 ng/µL of chloramphenicol with growth at 37°C to a density ($OD_{600\ nm}$) of 1.0. Bacterial Cultures were cooled over 1 hour to 18 degrees Celsius and recombinant protein expression was induced by addition of 1 mM IPTG and incubation at 18˚C for 18 hours. The cells were collected by centrifugation, and the bacterial cell pellets were resuspended in IMAC Lysis Buffer (50 mM HEPES, pH 7, 150 mM NaCl, 5 mM ß-mercaptoethanol, 10% [w/v] glycerol, 1 mM Benzamidine, 100 µM Leupeptin, 100 µM Antipain, 1 µM Pepstatin), flash frozen in liquid $N_2$, and stored at -80°C.

*Purification:* *E. coli* expressing GFP fusion proteins resuspended in IMAC Lysis Buffer were thawed on wet ice, diluted with an equal volume of IMAC Dilution Buffer (50 mM HEPES, pH 7, 1.85 M NaCl, 10 mM Imidazole, 5 mM ß-mercaptoethanol, 10% [w/v] glycerol, 1 mM Benzamidine, 100 µM Leupeptin, 100 µM Antipain, 1 µM Pepstatin), and lysed with multiple passages through an Avestin Emulsiflex-C5 high pressure homogenizer at ~10,000 PSI. Soluble bacterial lysate was isolated by centrifugation of cellular debris in a Beckman Avanti J-26 XPI centrifuge in a JA25.5 rotor at 4˚C for 40 minutes at 19,500 RPM. Clarified soluble bacterial lysate was applied to 15 mL of NiNTA resin (Qiagen) for 2 hours in batch with end-over-end mixing and washed in a glass Econo-Column (BioRad) with 500 mL of IMAC Wash Buffer A (50 mM HEPES, pH 7, 1 M NaCl, 25 mM Imidazole, 5 mM ß-mercaptoethanol, 10% [w/v] glycerol, 1 mM Benzamidine) and 100 mL of IMAC Wash Buffer B (50 mM HEPES, pH 7, 150 mM NaCl, 30 mM Imidazole, 5 mM ß-mercaptoethanol, 10% [w/v] glycerol, 1 mM Benzamidine). GFP fusion proteins were eluted from NiNTA resin with 100 mL of IMAC Elution Buffer (50 mM HEPES, pH 7, 150 mM NaCl, 300 mM Imidazole, 5 mM ß-mercaptoethanol, 10% [w/v] glycerol, 1 mM Benzamidine). NiNTA eluate was cleaved with Tobacco Etch Virus protease overnight (≥ 16 hours) at 4˚C, diluted with 9 volumes of Ion Exchange Buffer A (20 mM Tris•HCl, pH 8, 1 mM DTT), and applied to Source 15Q resin (GE Healthcare). Fusion proteins were eluted from the Source 15Q resin with 5-60% Ion Exchange Buffer B over 40 column volumes (20 mM Tris•HCl, pH 8, 1 M NaCl, 1 mM DTT). For GFP-TetR-$p300_{HAT}$, fractions containing the fusion protein as determined by SDS-PAGE with Coomassie Brilliant Blue Staining, were pooled, diluted with 9 volumes Ion Exchange Buffer A, applied to Source 15S resin (GE Healthcare), and eluted with 5-55% Ion Exchange Buffer B over 50 column volumes. GFP-TetR protein from Source 15Q cation exchange chromatography and GFP-TetR-p300 protein from Source 15S anion exchange chromatography were further purified by size exclusion chromatography using either a HiLoad SD200 26/60 pg column (GFP-TetR) or a Superdex 200 Increase 10/300 GL column (GFP-TetR-p300) in Gel Filtration Buffer (20 mM Tris•HCl, pH 8, 150 mM NaCl, 10% [w/v] glycerol, 1 mM DTT). The purest peak fractions of either GFP-TetR or GFP-TetR-$p300_{HAT}$, as determined by SDS-PAGE with Coomassie Brilliant Blue Staining, were pooled, concentrated, and quantified using absorbance at 485 nm and the known molar extinction coefficient for superfolder GFP of 83300/M•cm. Purified GFP-TetR and GFP-TetR-$p300_{HAT}$ were aliquoted, flash frozen with liquid $N_2$, and stored at -80˚C.

**Reconstitution of Histone Octamers**

Aliquots of histone H4 and H3 or histone H2A and fluorophore labeled or unlabeled H2B at 200 or 240 nmol quantities, respectively, were thawed on wet ice and diluted to 3 mL volume each in Histone Unfolding Buffer and combined for a final total volume of 12 mL. The histone mix was dialyzed in 1000 Dalton MWCO dialysis tubing (Spectra/Por) three times against 2 liters of Refolding Buffer (10 mM Tris•HCl, pH 7.5, 2 M NaCl, 1 mM EDTA, 5 mM ß-mercaptoethanol) with the second or third dialysis steps proceeding overnight. The dialysate was filtered by passage through a Whatman GD/XP 25 mm 0.45µm syringe filter. Refolded histone octamer was isolated by size exclusion chromatography of dialysate with a HiLoad SD200 26/60 pg column. Peak Fractions were analyzed by 15% PAGE-SDS analysis for stoichiometry of core histone proteins and those with clear histone octamers were pooled, concentrated by with a 10,000 Dalton MWCO centrifugal concentrator, and quantified by measuring absorbance at 280, 495, and 590 as well as absorbance at and the calculated molar extinction coefficients (https://web.expasy.org/protparam/ or Thermo Scientific) for histone octamer, AF488, and AF594 of 44700/M•cm, 73000/M•cm, and 92000/M•cm, respectively. 100% labeled histone octamers were confirmed by the presence of 2:1 stoichiometric excess of fluor to histone octamer. Purified and differentially labeled histone octamers were aliquoted, flash frozen with liquid $N_2$, and stored at -80 ˚C.

**Isolation and Purification of 12x601, 6x601, and 4x601 Array DNA.**

p12x601 was transformed into $dam^-$/$dcm^-$ *E. coli* strain ER2925 and plated onto LB agar plates supplemented with 20 ng/µL Zeocin for growth overnight. 6 liters of LB with 20 ng/µL of Zeocin were incubated with shaking overnight at 37 ˚C following their inoculation with an 8 hour liquid LB and 20 ng/µL Zeocin preculture started from a single colony on the LB agar plate. The 6 liters of turbid culture were harvested the next morning by centrifugation in a Sorvall RC3C Plus centrifuge, and plasmid DNA was purified using a Qiagen Plasmid Giga Kit according to the manufacturers instructions. 12x601 array DNA was cut from carrier plasmid DNA by incubation overnight (≥ 16 hours) with 5,000 units of EcoRV-HF restriction endonuclease (New England Biolabs) in 1x CutSmart Buffer (20 mM Tris•OAc, pH 7.9, 50 mM KOAc, 10 mM $Mg[OAc]_2$, 0.1 mg/mL BSA). Reaction was stopped by addition of 20 mM EDTA and DNA was purified by Phenol:Chloroform:Isoamyl Alcohol (25:24:1) Extraction and Ethanol Precipitation. DNA was resuspended in TE500 (10 mM Tris•HCl, pH 7.5, 1 mM EDTA, 500 mM NaCl) and 12x601 was size-fractionated by PEG precipitation with drop-by-drop addition of 30% PEG-8000 in TE500 with DNA solution under vortex. Before use in nucleosome assembly, >99% purity of 12x601 DNA array was confirmed by 1% agarose gel electrophoresis in 1xTAE (40 mM Tris•OAc, pH 8.6, 20 mM OAc, 1 mM EDTA) and staining with ethidium bromide (Bio-Rad). 6x601 or 4x601 array DNA were prepared by digesting 12x601 array DNA with SacI or NcoI and KpnI restriction endonucleases in 1x CutSmart Buffer. DNA was purified by Phenol:Chloroform:Isoamyl Alcohol (25:24:1) Extraction and Ethanol Precipitation. >99% purity of 6x601 or 4x601 DNA array was confirmed by 1% agarose gel electrophoresis in 1xTAE (40 mM Tris•OAc, pH 8.6, 20 mM OAc, 1 mM EDTA) and staining with ethidium bromide (Bio-Rad).

**Preparation of Ligation-competent 601 DNA**

98x102 µL polymerase chain reactions (20 mM Tris•HCl, pH 8.8, 10 mM $[NH_4]2SO_4$, 10 mM KCl, 2 mM $MgSO_4$, 0.1% Triton®-X-100, 500 nM oligonucleotides, 25 pg/µL 601 template, 25 mU/µL TAQ DNA polymerase [NEB], 250 µM each dNTPs, 1% DMSO) were performed with (1) 2 minutes at 95˚C, (2) 40 cycles of 30 seconds at 95˚C, 30 seconds at 60˚C, and 1 minute at 72˚C, (3) 2 minutes at 72˚C, and (4) cooling to 4˚C. Separate reactions were pooled in a 3,000 Dalton MWCO centrifugal concentrator (Amicon) and the 10 mL PCR reaction was stopped by addition of 400 µL of 0.5 M EDTA. Unincorporated dNTPS and oligonucleotides were removed, and PCR amplified DNA was washed by repeated centrifugal concentration and subsequent dilution of PCR reaction with 1xTE (10 mM Tris•HCl, pH 7.5, 1 mM EDTA). Concentrated and 1xTE-washed PCR product was purified by Phenol:Chloroform:Isoamyl Alcohol (25:24:1) Extraction and Ethanol Precipitation. Amplified DNA was digested with 1,000 units of restriction endonuclease BstXI (NEB) overnight (≥ 16 hours). Cut PCR Product was purified by Phenol:Chloroform:Isoamyl Alcohol (25:24:1) Extraction and Ethanol Precipitation and resuspended in TE50 (10 mM Tris•HCl, pH 7.5, 1 mM EDTA, 50 mM NaCl). Ligation-competent dsDNA was isolated by size exclusion chromatography of BstXI-digested DNA in TE50 using a Superdex 200 Increase 10/300GL column and concentrated for nucleosome assembly using 3,000 Dalton MWCO centrifugal concentrator (Amicon).

**Preparation of Polynucleosomal Arrays and Mononucleosomes**

***Setup of Nucleosomal Assemblies.*** Quantified DNA template and histone octamers of choice were thawed on wet ice. 601 sequence-containing DNA dissolved in 1xTE and an equal volume of 4M Assembly Buffer (10 mM Tris•HCl, pH 7.5, 1 mM EDTA, 4 M NaCl, 2 mM DTT) were mixed thoroughly on on ice, followed by addition equimolar quantities of histone octamer ratio relative to 601 nucleosome positioning sequences in the template. Final concentrations of octamer/601 varied between 1-5 µM, and fluorophor labeled octamer varied between 1 and 100% labeled nucleosomes with no detectable difference in assembly efficiency. Assembly of mutant *X. laevis* histone octamers into nucleosomes was aided by addition of 0.2 molar excess labeled histone H2A/H2B dimers. Histone octamers and 601-containing DNA templates were moved into <8,000 Dalton MWCO dialysis chambers equilibrated in High Salt Assembly Buffer (10 mM Tris•HCl, pH 7.5, 1 mM EDTA, 2 M KCl, 1 mM DTT).

***Salt Dialysis-mediated Assembly of Nucleosomes.*** First, dialysis chambers were placed in 2 L of High Salt Assembly Buffer and salt concentration was lowered by continuous dilution with 2 L of Low Salt Assembly Buffer (10 mM Tris•HCl, pH 7.5, 1 mM EDTA, 200 mM KCl, 1 mM DTT) using a peristaltic pump at 0.8 mL/min and vigorous stirring at 4˚C. Second, after exhaustion of Low Salt Assembly Buffer, the dialyzing volume was reduced to 500 mL of liquid and the salt concentration was lowered further by continuous dilution with 1 L of No Salt Assembly Buffer (10 mM Tris•HCl, pH 7.5, 1 mM EDTA, 1 mM DTT) at 0.8 mL/min at 4˚C with constant vigorous stirring. Last, the dialysis chambers were dialyzed against No Salt Assembly Buffer for at least for hours at 4˚C.

***Sucrose Gradient-mediated Purification of Nucleosomes.*** Following salt-mediated dialysis, assembled polynucleosomal arrays or mononucleosomes in No Salt Assembly Buffer were applied to linear 15-40% or 5-20% sucrose gradients in No Salt Assembly Buffer. Sucrose gradient fractions containing assembled nucleosomes were concentrated in 10,000 Dalton MWCO centrifugal concentrators (Amicon).

***Quantitation of Nucleosome Concentration.*** To quantitate final chromatin concentrations, 1 µL of assembled polynucleosomal arrays or mononucleosomes or their DNA templates of known DNA concentration were added to 99 µL of SDS/PK Buffer (45 mM Tris•HCl, pH 7.5, 9 mM EDTA, 1% SDS) and incubated for 30 minutes at room temperature. DNA was purified by either Phenol:Chloroform:Isoamyl alcohol extraction and ethanol precipitation or processing samples with a Qiagen PCR purification Kit. The quantity of DNA in assembled mononucleosomes was determined according to a standard curve generated by serial dilution of DNA purified from unassembled template.

***Quality Assurance of Nucleosomal Assembly.*** Mononucleosomal assemblies were assessed for quality by electrophoretic mobility shift assay on a 2% agarose gel. Polynucleosomal assemblies were assessed for quality by differential digestion relative to purified unassembled 12x601 DNA with either EcoRI-HF (linker DNA sites) or BsiWI-HF (nucleosomal DNA sites) in 1x CutSmart Buffer. DNA fragments were purified with a Qiagen PCR purification Kit and analyzed on a 1% Agarose gel in 1xTAE.

**Preparation of 384-well Microscopy Plates**

***mPEGylation of Silica.*** 384-well microscopy plates (Brooks Life Science Systems Matriplate) were washed with 5% Hellmanex at 65˚C for 4 hours and then rinsed copiously with ≥ 18 MΩ $H_2O$. Silica was etched with 1 M NaOH for 1 hour at room temperature and then rinsed copiously with ≥ 18 MΩ $H_2O$. Depolymerized Silica was covalently bonded overnight ( ≥ 18 hours) at room temperature to 25 mg/mL 5K mPEG-silane (PEGWorks) suspended in 95% Ethanol. Plate was washed once with 95% Ethanol, rinsed with copious amounts of ≥ 18 MΩ $H_2O$, and completely dried in a chemical hood over 3-4 hours. PEGylated microscopy plate was sealed until individual wells' use with an adhesive PCR plate foil (Thermo).

***Passivation of Well with Bovine Serum Albumin.*** Following PEGylation, foil was cut above individual wells prior to their use and both plastic and PEGylated glass were passivated by incubation with freshly prepared 100 mg/mL BSA for 30 minutes. Wells were rinsed once with ≥ 18 MΩ $H_2O$ to remove excess BSA, and microscopy samples (15-40 µL in volume) were immediately added to the empty well. Desiccation of microscopy samples was limited following their addition to the plate by sealing with transparent scotch tape.

**Phase Separation and Microscopy of Polynucleosomal Arrays**

***Phase Separation of Polynucleosomal Arrays.*** Nucleosomal arrays, with a 1 in 100 histone H2B proteins labeled with a fluorophore, was first equilibrated in Chromatin Dilution Buffer (25 mM Tris•OAc, pH 7.5, 5 mM DTT, 0.1 mM EDTA, 0.1 mg/mL BSA, 5%[w/v] glycerol) and incubated for 5 minutes at room temperature. Phase separation was induced, unless otherwised indicated, by addition of 1 volume of Phase Separation Buffer (25 mM Tris•OAc, pH 7.5, 0.1 mM EDTA, 5 mM DTT, 0.1 mg/mL BSA, 5%[w/v] glycerol, 300 mM KOAc, 1 mM Mg[OAc]$_2$, 2 µg/mL Glucose Oxidase [SIGMA cat. no. G2133], 350 ng/mL Catalase [SIGMA cat. no. C1345], 4 mM Glucose) to 750 nM chromatin equilibrated in Chromatin Dilution Buffer. 30 minutes after addition of Phase Separation Buffer reactions were gently mixed and added to the well of a PEGylated and BSA passivated microscopy plate.

***In-phase Chromatin Binding and Microscopy.***

Histone H1: Following a 30 minute incubation period after addition of Phase Separation Buffer, 10x concentrated histone H1 in Histone H1 Buffer A or Histone H1 Buffer A alone was added to phase separated chromatin at a 1:1 stoichiometric ratio of histone H1:nucleosomes and samples were immediately added to a PEGylated and BSA passivated microscopy plate. Spinning disk confocal microscopy images were acquired after 1 hour of incubation in the 384-well microscopy plate.

GFP Fusion Proteins: Following a 30 minute incubation period after addition of Phase Separation Buffer either supplemented with or without 2 µg/mL Doxycycline, 4 µM GFP-TetR or GFP-TetR-p300$_{HAT}$ diluted in Gel Filtration Buffer was added in 0.333 reaction volumes to phase-separated chromatin and incubated for 30 minutes at room temperature. Samples were then moved to a PEGylated and BSA passivated microscopy plate. Spinning disk confocal microscopy images were acquired after 1 hour of incubation in the 384-well microscopy plate. For time-resolved microscopy of GFP-TetR-p300$_{HAT}$-dependent histone acetylation and droplet dissolution, 1 µL of 10 mM AcetylCoA was added to the microscopy well and allowed to mix by diffusion.

***Confocal Fluorescence Microscopy.*** Confocal fluorescence microscopy images were captured on a Nikon Eclipse Ti microscope base equipped with a Yokogawa CSU-X1 spinning disk confocal scanner unit, 100 X 1.49 NA objective, and Andor EM-CCD camera. Fluorescence Recovery After Photobleaching (FRAP) was achieved with a TIRF/iLAS2 FRAP Module (Biovision) and Rapp UGA-40 Phototargeter.

***Microscopy Data Analysis.*** Microscope Images were analyzed with ImageJ (*6*). Unless otherwise described, equivalent brightness and contrast were used when depicting microscopy images in a given panel. Microscopy data processed by ImageJ was graphed using the R Statistical Package (*7*).

**In Vitro Histone Acetylation**

12x601 nucleosomal arrays assembled with wild-type X. laevis histone octamers labeled with Atto565 fluorophores or basic-patch mutant X. laevis histone octamers labeled with Cy5 fluorophores were equilibrated in Chromatin Dilution Buffer (25 mM Tris•OAc, pH 7.5, 5 mM DTT, 0.1 mM EDTA, 0.1 mg/mL BSA, 5% [w/v] glycerol) and incubated for 5 minutes at room temperature. Phase separation was induced by addition of 1 volume of Phase Separation Buffer (25 mM Tris•OAc, pH 7.5, 0.1 mM EDTA, 5 mM DTT, 0.1 mg/mL BSA, 5% [w/v] glycerol, 300 mM KOAc, 1 mM Mg[OAc]$_2$, 2 µg/mL Glucose Oxidase [SIGMA cat. no. G2133], 350 ng/mL Catalase [SIGMA cat. no. C1345], 4 mM Glucose, ± 2 μg/mL Doxycycline) to chromatin (1μM nucleosomes) equilibrated in Chromatin Dilution Buffer. 30 minutes after addition of Phase Separation Buffer, GFP-TetR-$p300_{HAT}$ was added at a final concentration of 250 nM and incubated for 30 minutes at room temperature. In vitro acetylation was triggered by addition of 400 μM AcetylCoA and reactions proceeded for 30 minutes at room temperature before stopping by addition of 1 reaction volume of 2x SDS-PAGE Sample Buffer (65.8 mM Tris•HCl, pH 6.8, 0.71 mM ß-Mercaptoethanol, 26.3% [w/v] glycerol, 2.1% [w/v] SDS, 0.01% bromophenol blue)

Reaction products from the in vitro acetylation reaction were run on a 15% PAGE-SDS gel and transferred to a PVDF membrane for analysis by western blotting. The PVDF membrane was blocked with 5% Milk in TBST (20 mM Tris•HCl, pH 7.4, 150 mM NaCl, 0.05% Tween) and blotted with a 1:5,000 dilution of Rabbit polyclonal antibody against histone H3K27 acetylation (Abcam ab4729) in 1% milk in TBST overnight at 4˚C. Primary antibody was washed off of the membrane with multiple washes of TBST and 1:10,000 mouse anti-rabbit HRP was incubated with the membrane for 1 hour at room temperature in 1% milk in TBST. Excess secondary antibody was washed from the membrane and the western blot signal was developed with Millipore Immobilon HRP substrate using a ChemiDoc Imaging System (Bio-Rad).

**Trypsinization of Core Histone Tails**

Sequencing grade Trypsin (Promega) was solubilized at a concentration of 100 ng/μL in 50 mM Acetic Acid, flash frozen, and stored at -80˚C prior to use in trypsinization reactions. Just prior to proteolysis, Trypsin was diluted in 9 volumes of 25 mM Tris•HCl, pH 8 to neutralize 50 mM Acetic Acid. Nucleosomes were trypsinized in Trypsin Digestion Buffer (25 mM Tris•OAc, pH 7.5, 0.1 mM EDTA, 5 mM DTT, 5% [w/v] glycerol, 0.25 ng/μL Trypsin) for 30 minutes at room temperature and reactions were stopped by addition of Aprotinin and BSA at a final concentration of 50 ng/μL and 0.25 mg/mL, respectively. Digestion was confirmed by 15% PAGE-SDS and Coomassie Brilliant Blue staining of core histone proteins and stopped reactions were added to phase separation assays exactly as described above.

**Ligation-dependent Assembly of Nucleosomal Arrays from Mononucleosomes.**

1 μM mononucleosomes with different linker DNA lengths were assembled with BstXI-digested DNA were ligated in Ligation Assembly Buffer (50 mM Tris•OAc, pH 7.5, 5 mM DTT, 200 μM ATP, 3 mM $Mg[OAc]_2$, 100 mM KOAc, 0.1 mg/mL BSA, 5% [w/v] glycerol, 1 U/μL T4 DNA Ligase [Enzymatics]) for 30 minutes at room temperature, and reactions were stopped by addition of 0.2 reaction volumes of 6x Reaction Stop Buffer (60 mM EDTA, pH 8, 400 mM KOAc). After quenching ligation by chelation of free magnesium, ligation reactions were moved into the well of an mPEGylated and BSA passivated microscopy plate. DNA was extracted by addition of SDS/PK buffer to 100 μL final volume and purification with a PCR Purification Kit (Qiagen) according to manufacturer's instructions. BsiWI-HF digestion of 601 DNA sequences within ligation products was used as a quantitative measure of the extent of ligation by T4 DNA Ligase.

**Absolute Quantitation of Nucleosome Concentration in Condensates and In Solution**

***In-phase Quantitation.*** Unlabeled and 8.3% (1:12) AF594-labeled 12x601 polynucleosomal arrays were assembled using the salt dialysis method and sucrose gradient purification as described above. Nucleosomal arrays were combined at to achieve 0.1% AF594- or AF488-labeled octamers and phase separation of chromatin was triggered as described above, but omitting $Mg(OAc)_2$ in the Phase Separation Buffer. Droplets with or without addition of equimolar histone H1 relative to nucleosomes were moved into an mPEGylated and BSA passivated microscopy well and mean fluorophore intensity within droplets was measured after 1 hour of incubation relative to a standard curve of free AF594 or AF488 dye.

***In-solution Quantitation.*** 12x601 polynucleosomal arrays were assembled with 100% AF594-labeled histone octamers using the salt dialysis method and sucrose gradient purification as described above. Phase separated was triggered exactly as described above and droplets were pelleting by 10 x 1 minute at maximum speed in a microcentrifuge. After equilibration for 10 minutes at room temperature supernatant above the droplets was moved into an mPEGylated and BSA passivated microscopy well and fluorophore intensity was measured relative to a standard curve of free AF594 dye.

**Electrophoretic Mobility Shift Assay**

Glycerol and NaCl were added to purified DNA or mononucleosomes at final concentrations of 10% w/v and 5 mM, respectively, and loaded into the wells of 6% polyacrylamide gels buffered with 0.5x TAE. Nucleic acid-containing samples were separated by electrophoresis at 100 volts for 40 minutes before imaging fluorescence of labeled nucleosomes with a ChemiDoc Imaging Station (BioRad) and staining of DNA using ethidium bromide.

***Primers for PCR Amplification of Ligation-competent 601 DNA.***

Primers used to amplify 601 DNA competent for ligation-depedendent assembly of polynucleosomal arrays from mononucleosomes *(listed 5' to 3').*

172NRL_Fwd: GATATCccacgcatatggATGTAACTGGAGAATCCCGGTGC
172NRL_Rev: GGATCCccagatgcatggGATGGGAACAGGATGTATATATCTGACACG

**3) <u>Supplementary References</u>**

1. E. Balleza, J. M. Kim, P. Cluzel, Systematic characterization of maturation time of fluorescent proteins in living cells. *Nature methods* **15**, 47-51 (2018).
2. A. G. Larson *et al.*, Liquid droplet formation by HP1alpha suggests a role for phase separation in heterochromatin. *Nature* **547**, 236-240 (2017).
3. K. Luger, T. J. Rechsteiner, T. J. Richmond, Expression and purification of recombinant histones and nucleosome reconstitution. *Methods in molecular biology* **119**, 1-16 (1999).
4. M. D. Wilson *et al.*, The structural basis of modified nucleosome recognition by 53BP1. *Nature* **536**, 100-103 (2016).
5. H. Klinker, C. Haas, N. Harrer, P. B. Becker, F. Mueller-Planitz, Rapid purification of recombinant histones. *PloS one* **9**, e104029 (2014).
6. C. A. Schneider, W. S. Rasband, K. W. Eliceiri, NIH Image to ImageJ: 25 years of image analysis. *Nature methods* **9**, 671-675 (2012).
7. R. C. Team, R: A language and environment for statistical computing., (2013).
8. A. R. Quinlan, I. M. Hall, BEDTools: a flexible suite of utilities for comparing genomic features. *Bioinformatics* **26**, 841-842 (2010).
